# Synaptonemal complex proteins direct and constrain the localization of crossover-promoting proteins during *Caenorhabditis elegans* meiosis

**DOI:** 10.1101/523605

**Authors:** Cori K. Cahoon, Jacquellyn M. Helm, Diana E. Libuda

**Author notes:** Corresponding Author Information: Diana E. Libuda University of Oregon Institute of Molecular Biology 1229 Franklin Blvd Eugene, OR 97403.

## Abstract

Crossovers (COs) between homologous chromosomes are critical for meiotic chromosome segregation and form in the context of the synaptonemal complex (SC), a meiosis-specific structure that assembles between aligned homologs. During *Caenorhabditis elegans* meiosis, central region components of the SC (SYP proteins) are essential to repair double-strand DNA breaks (DSBs) as COs, but the roles of these SYP proteins in promoting CO formation are poorly understood. Here, we investigate the relationships between the SYP proteins and conserved CO-promoting factors by examining the immunolocalization of these factors in meiotic mutants where SYP proteins are absent, reduced, or mis-localized. Although COs do not form in *syp* null mutants, CO-promoting proteins COSA-1, MSH-5, and ZHP-3 nevertheless become co-localized at a variable number of DSB-dependent sites during late prophase, reflecting an inherent affinity of these factors for DSB repair sites. In contrast, in mutants where SYP proteins are present but form aggregates or display abnormal synapsis, CO-promoting proteins consistently track with SYP-1 localization. Moreover, CO-promoting proteins usually localize to a single site per SYP-1 structure, even in SYP aggregates or in mutants where SC forms between sister-chromatids, suggesting that CO regulation occurs within these structures. Further, we find that sister chromatids in the meiotic cohesin mutant *rec-8* require both CO-promoting proteins and the SC to remain connected. Taken together, our findings support a model in which SYP proteins promote CO formation by directing and constraining the localization of CO-promoting factors to ensure that CO maturation occurs only between properly aligned homologous chromosomes.

**Article Summary:** Errors during meiosis are the leading cause of birth defects and miscarriages in humans. Thus, the coordinated control of meiosis events is critical for the faithful inheritance of the genome each generation. The synaptonemal complex (SC) is a meiosis-specific structure that assembles between homologs chromosomes and is critical for the establishment and regulation of crossovers, which ensure the accurate segregation of the homologous chromosomes at meiosis I. Here we show that the SC proteins function to regulate crossovers by directing and constraining the localization of proteins involved in promoting the formation of crossovers.

## Introduction

During sexual reproduction, generation of haploid gametes from diploid germ cells involves substantial reorganization of chromosomes within the nucleus and formation of specialized meiosis-specific chromosome structures. In preparation for segregating to opposite poles at the meiosis I division, homologous chromosome pairs align along their full lengths and assemble a structure known as the synaptonemal complex (SC) between them. The SC structure is composed of axial elements that assemble along the lengths of conjoined pairs of sister chromatids (known as lateral elements (LEs) in the context of assembled SC), and a set of proteins that comprise the central region (CR) of the SC that link the parallel-aligned homolog axes (reviewed in Cahoon and hawley 2016). Multiple SC-CR proteins function together to span the distance between the LEs and are required for normal meiosis all organisms that assemble the SC. Although, the collective properties of the SC-CR and its constituent proteins and the way these proteins contribute to a successful outcome of meiosis remains a subject of active investigation.

In the current work, we investigate the function of the SC-CR proteins during meiosis in the nematode *Caenorhabditis elegans*. Four different components of the SC-CR have been identified in *C. elegans*, termed SYP-1, SYP-2, SYP-3 and SYP-4 (<u>sy</u>naptonemal complex <u>p</u>rotein). These proteins localize between the LEs of the SC and are interdependent for localization and stability (Schild-prufert *et al.* 2011). analysis of *syp* mutants has demonstrated that the syp proteins are required both to stabilize homolog pairing between homologous chromosomes and to promote the formation of crossover (co) recombination events (macqueen *et al.* 2002; Colaiacovo *et al.* 2003; Smolikov *et al.* 2007; Smolikov *et al.* 2009). The SYP proteins also play a role in limiting the number of COs that form during meiosis, as partial depletion of SYP proteins by RNAi results in an increase in the number of chromosomes with more than one CO and an attenuation of CO interference (Libuda *et al.* 2013). Further, recent studies have provided evidence for liquid crystalline-like behavior of the SC-CR and have revealed dynamic properties of the SC-CR that change during the course of meiotic prophase progression (Rog and dernburg 2015; Mlynarczyk-evans and villeneuve 2017; Pattabiraman *et al.* 2017; Rog *et al.* 2017). Moreover, studies have identified properties of the SC-CR, including phosphorylation state of its components and recruitment of polo-like kinase PLK-2 by CHK-2, that are altered in response to formation of CO recombination intermediates (Kim *et al.* 2015; Nadarajan *et al.* 2016; Nadarajan *et al.* 2017; Pattabiraman *et al.* 2017). Despite these advances in understanding, however, what the SC-CR proteins are doing to promote the formation of interhomolog COs during meiosis remains poorly understood.

In the context of an assembled SC, a set of CO-promoting proteins (MSH-5, COSA-1, ZHP-1, ZHP-2, ZHP-3, and ZHP-4) are loaded on chromosomes during *C. elegans* meiosis to promote and license the repair of these double strand DNA breaks (DSBs) as interhomolog COs. Following the formation of DSBs by the conserved endonuclease SPO-11, the CO-promoting proteins MSH-5 (a component of the meiosis-specific MutSγ complex) and COSA-1 (a cyclin-related protein specific to metazoan meiosis) form multiple DSB-dependent foci in early pachytene prior to reducing down in number in late pachytene, marking the 6 CO sites in *C. elegans* meiosis (Kelly *et al.* 2000; Yokoo *et al.* 2012). In contrast, ZHP-1, ZHP-2, ZHP-3, ZHP-4 (RING domain containing proteins) coat the SC before localizing into distinct foci that colocalize with the 6 CO sites marked by MSH-5 and COSA-1 in late pachytene (Jantsch *et al.* 2004; Bhalla *et al.* 2008; Nguyen *et al.* 2018; Zhang *et al.* 2018). Although analysis of null mutants of the CO-promoting proteins demonstrates that these proteins are interdependent for their localization and are required for CO formation, the mechanism of how the CO-promoting proteins function together to establish a CO is unknown.

Recent evidence indicates that the SC proteins envelop CO-designated sites marked by the CO-promoting proteins (Woglar and villeneuve 2018); however, the relationship between the CO-promoting proteins and the SC is still largely unclear.

Here we address how the SC-CR protein promote formation of interhomolog COs by investigating how the SC central region proteins contribute to localization of conserved CO-promoting factors that normally become localized at CO sites. Our data suggest that SYP proteins direct and constrain where CO-promoting factors can localize, thereby ensuring that COs form appropriately, *i.e.* between the DNA molecules of successfully paired homologous chromosomes.

## Materials and Methods

### C. elegans strains, genetics, and culture conditions

All strains are from the Bristol N2 background and were maintained and crossed at 20**°**C under standard conditions. Temperatures used for specific experiments are indicated below. For all experiments with meiotic mutants, homozygous mutant worms were derived from balanced heterozygous parents by selecting progeny lacking a dominant marker (Unc and/or GFP) associated with the balancer.

The following strains were used in this study: N2: Bristol wild-type strain.

AV198: *spo-11(ok79) IV; syp-1(me17) V / nT1[unc-?(n754) let-? qIs50] (IV;V)*.

AV276: *syp-2(ok307) V / nT1[unc-?(n754) let-?(m435)] (IV;V)*.

AV278: *spo-11 IV; syp-2(ok307) V / nT1[unc-?(n754) let-? qIs50] (IV;V)*.

AV307: *syp-1(me17) V / nT1[unc-?(n754) let-? qIs50] (IV;V)*.

AV596: *cosa-1(tm3298)/ qC1[qIs26] (III)*.

AV630: *meIs8[unc-119(+) Ppie-1::gfp::cosa-1] II*.

AV647: *meIs8[unc-119(+) Ppie-1::gfp::cosa-1] II; spo-11(me44) IV / nT1[unc-?(n754) let-? qIs50] (IV;V)*.

AV671: *meIs8[unc-119(+) Ppie-1::gfp::cosa-1] II; him-3(e1256) IV*.

AV686: *meIs8[unc-119(+) Ppie-1::gfp::cosa-1] II; rec-8(ok978) IV / nT1[qIs51] (IV;V)*.

AV687: *syp-3(ok758) I / hT2[bli-4(e937) let-?(q758) qIs48] (I;III); meIs8[unc-119(+) Ppie-1::gfp::cosa-1] II*.

AV688: *meIs8[unc-119(+) Ppie-1::gfp::cosa-1] II; syp-2(ok307) V / nT1[unc-?(n754) let-?(m435)] (IV;V)*.

AV689: *meIs8[unc-119(+) Ppie-1::gfp::cosa-1] II; him-3(gk149) IV / nT1[qIs51] (IV;V)*.

AV695: *meIs8[unc-119(+) Ppie-1::gfp::cosa-1] II; mnT12 (X;IV)*.

AV697: *meIs8[unc-119(+) Ppie-1::gfp::cosa-1] II; htp-3(y428) ccIs4251 I / hT2[bli-4(e937) let-?(q782) qIs48] (I,III)*.

AV699: *meIs8[unc-119(+) Ppie-1::gfp::cosa-1] II; syp-1(me17) V / nT1[unc-?(n754) let-? qIs50] (IV;V)*.

AV700: *him-3(gk149) IV / nT1[qIs51] (IV;V); syp-2(ok307) V / nT1[qIs51] (IV;V)*.

CB1256: *him-3(e1256) IV*.

CV2: *syp-3(ok758) I / hT2[bli-4(e937) let-?(q758) qIs48] (I;III).* TY4986: *htp-3(y428) ccIs4251 I / hT2[bli-4(e937) let-?(q782) qIs48] (I,III).* VC418: *him-3(gk149) IV / nT1[qIs51] (IV;V)*.

VC666: *rec-8(ok978) IV / nT1[qIs51] (IV;V)*.

DLW1: *cosa-1(tm3298)/ qC1[qIs26] III; rec-8(ok978) IV/nT1 [qIs51] (IV;V)*.

DLW12: *GFP::COSA-1 II; rec-8 (ok978)/ nT1 [qIs51] (IV;V)IV; syp-2(ok307) V/nT1 (IV;V)*.

Additional information on strains:

*qIs48* contains *[Pmyo-2::gfp; Ppes-10::gfp; Pges-1::gfp].*

*qIs50* contains *[Pmyo-2::gfp; Ppes-10::gfp; PF22B7.9::gfp].*

*qIs51* contains *[Pmyo-2::gfp; Ppes-10::gfp; PF22B7.9::gfp]*.

**syp-1 *partial depletion by RNAi.*** Partial depletion of *syp-1* by RNAi was performed as in (Libuda *et al.* 2013). Worms were synchronized at the L1 phase by bleaching adults and allowing resultant eggs to hatch on unseeded NGM plates at 20**°**C for 20-24 hrs.

Synchronized L1s were then washed off of the unseeded NGM plates with M9 and placed on NGM+IPTG+Amp plates that were poured within 30 days of use and freshly seeded one day before use with *Escherichia coli* HT115 cells containing either a fragment of the *syp-1/F26D2.2* gene in the L4440 vector (Ahringer Lab RNAi library) or, the empty vector (referred to as “control RNAi” in figures and text). The RNAi plates with L1s were then placed at 25**°**C for 40-48 hrs and then their gonads dissected for immunofluorescence.

### Immunofluorescence

Immunofluorescence was performed as in (Libuda *et al.* 2013). Gonads from adult worms at 18-24 hours post-L4 stage were dissected in 1x egg buffer with 0.1% Tween on VWR Superfrost Plus slides, fixed for 5 min in 1% paraformaldehyde, flash frozen with liquid nitrogen, and then fixed for 1 minute in 100% methanol at −20**°**C. Slides were washed 3 x 5 min in 1x PBST and blocked for one hour in 0.7% BSA in 1x PBST. Primary antibody dilutions were made in 1x PBST and added to slides. Slides were covered with a parafilm coverslip and incubated in a humid chamber overnight (14-18 hrs). Slides were washed 3 x 10 min in 1x PBST. Secondary antibody dilutions were made at 1:200 in 1x PBST using Invitrogen goat or donkey AlexaFluor labeled antibodies and added to slides. Slides were covered with a parafilm coverslip and placed in a humid chamber in the dark for 2 hrs. Slides were washed 3 x 10 min in 1x PBST in the dark. All washes and incubations were performed at room temperature, unless otherwise noted. 2 μg/ml DAPI was added to slides and slides were subsequently incubated in the dark with a parafilm coverslip in a humid chamber. Slides were washed once for 5 min in 1x PBST prior to mounting with Vectashield and a 20 x 40 mm coverslip with a 170 ± 5 μm thickness. Slides were sealed with nail polish immediately following mounting and then stored at 4**°**C prior to imaging. For structured illumination microscopy imaging (SIM), slides were made as described above with the following modification. All SIM slides were mounted in Prolong Gold (ThermoFisher, P36930) and left to harden at room temperature for 2-3 days prior to imaging. All slides were imaged (as described below) within two weeks of preparation. The following primary antibody dilutions were used: rabbit anti-GFP (1:1000) (Yokoo *et al.* 2012); chicken anti-GFP (1:1000) (Abcam 13970); guinea pig anti-ZHP-3 (1:500) (Bhalla *et al.* 2008); rabbit anti-MSH-5 (1:10000) (Novus #3875.00.02); guinea pig anti-SYP-1 (1:200) (Macqueen *et al.* 2002); goat anti-SYP-1 (1:1500) (Harper *et al.* 2011) and, chicken anti-HTP-3 (1:500) (Macqueen *et al.* 2005).

### Imaging

Immunofluorescence slides were imaged at 512 x 512 pixel dimensions on an Applied Precision DeltaVision microscope with a 63x lens with 1.5x optivar. Images were acquired as Z-stacks at 0.2 μm intervals and deconvolved with Applied Precision softWoRx deconvolution software. For quantification of GFP::COSA-1 foci, nuclei that were in the last 4-5 rows of late pachytene and were completely contained within the image stack were analyzed. Foci were quantified manually from deconvolved three-dimensional stacks. For quantification of chiasmata and visualization of chiasmata, individual chromosomes from a single diakinesis nucleus were cropped and rotated in three dimensions using Volocity three-dimensional rendering software. Images shown are projections through three-dimensional data stacks encompassing whole nuclei, generated with a maximum-intensity algorithm with the softWoRx software.

For SIM, slides were imaged at 2430 x 2430 pixel dimensions on a Zeiss ELYRA S.1 / LSM 880 microscope with a Plan Apochromat 63x (1.4 NA) oil lens. Images were acquired as a Z-stack at 0.110 µm interval with 3 rotations and were processed using the Zeiss ZEN software for both SIM reconstruction and channel alignment (alignment calibration based off 100-nm TetraSpeck beads from ThermoFisher). Maximum-intensity projections were generated using FIJI (NIH). Images were adjusted for brightness and contrast.

### Statistics

All p-values reported are two-tailed and calculated from Mann-Whitney tests, which are robust non-parametric statistical tests appropriate for the relevant data sets.

## Data Availability

All strains are available upon request. Figure S1A shows localization of COSA-1 and MSH-5 in *him-3, htp-3* and *rec-8* mutants. Figure S1B shows localization of COSA-1 and ZHP-3 in *rec-8*. Figure S2 shows the localization of RAD-51 in *rec-8* and *cosa-1; rec-8* mutants. Supplemental materials available at Figshare.

## Results and Discussion

### CO-promoting proteins co-localize to DSB-dependent events in late pachytene of mutants lacking synaptonemal complex central region proteins

Crossing over between homologous chromosomes during *C. elegans* meiosis is dependent on the SYP proteins, encoded by the *syp-1, syp-2, syp-3 and syp-4* genes. Initiation of meiotic recombination by formation of DSBs occurs in *syp* null mutants, but repair of these breaks does not yield interhomolog COs (Macqueen *et al.* 2002; Colaiacovo *et al.* 2003; Smolikov *et al.* 2007; Smolikov *et al.* 2009). However, immunolocalization of several conserved CO-promoting proteins (COSA-1, MSH-5 and ZHP-3) revealed that these pro-CO proteins nevertheless eventually become concentrated together at several sites per nucleus during late prophase in mutants lacking the SYP proteins (Figure 1A). In whole-mount preparations of wild-type gonads, ZHP-3 (an E3 ligase) is localized along the lengths of the chromosomes in nuclei at the mid-pachytene stage of meiotic prophase, and MSH-5 is detected as foci in excess of the eventual number of COs at this stage (Jantsch *et al.* 2004; Bhalla *et al.* 2008; Zhang *et al.* 2018). Upon transition to the late pachytene stage, COSA-1 is detected as bright foci at nascent CO sites, colocalizing with MSH-5, and ZHP-3 tracks gradually reduce and retract toward the COSA-1 foci (Yokoo *et al.* 2012). In *syp* null mutants, nuclei in the region of the gonad that would correspond to mid pachytene in wild-type gonads exhibit prolonged persistence of chromosome clustering (Macqueen *et al.* 2002; Colaiacovo *et al.* 2003; Smolikov *et al.* 2007; Smolikov *et al.* 2009). ZHP-3 is not detected along the lengths of the chromosomes in these nuclei, this result is expected based on previous reports that ZHP-3 localization is SYP-dependent (Jantsch *et al.* 2004). Notably, we found that upon eventual release from chromosome clustering and transition to a late pachytene-like dispersed chromosome organization, ZHP-3 does eventually form foci, and most of these foci colocalize with MSH-5 or COSA-1 (Figure 1A). The ZHP-3, MSH-5 and COSA-1 foci observed in the *syp* mutants are of weaker intensity than those observed in wild-type or other mutant situations where SYP proteins are present (see below). This decrease in intensity suggests that the CO-promoting proteins may have a weak affinity for such repair intermediates, which is both directed and strengthened by the presence of SYPs between homologs. Further, in contrast to the highly reproducible number of COSA-1 foci (6 per nucleus) detected in wild-type, numbers of COSA-1 foci observed in *syp* mutants are reduced and statistically different from wild-type (Figure 1B; *P<*0.0001, Mann Whitney).

**Figure 1.**
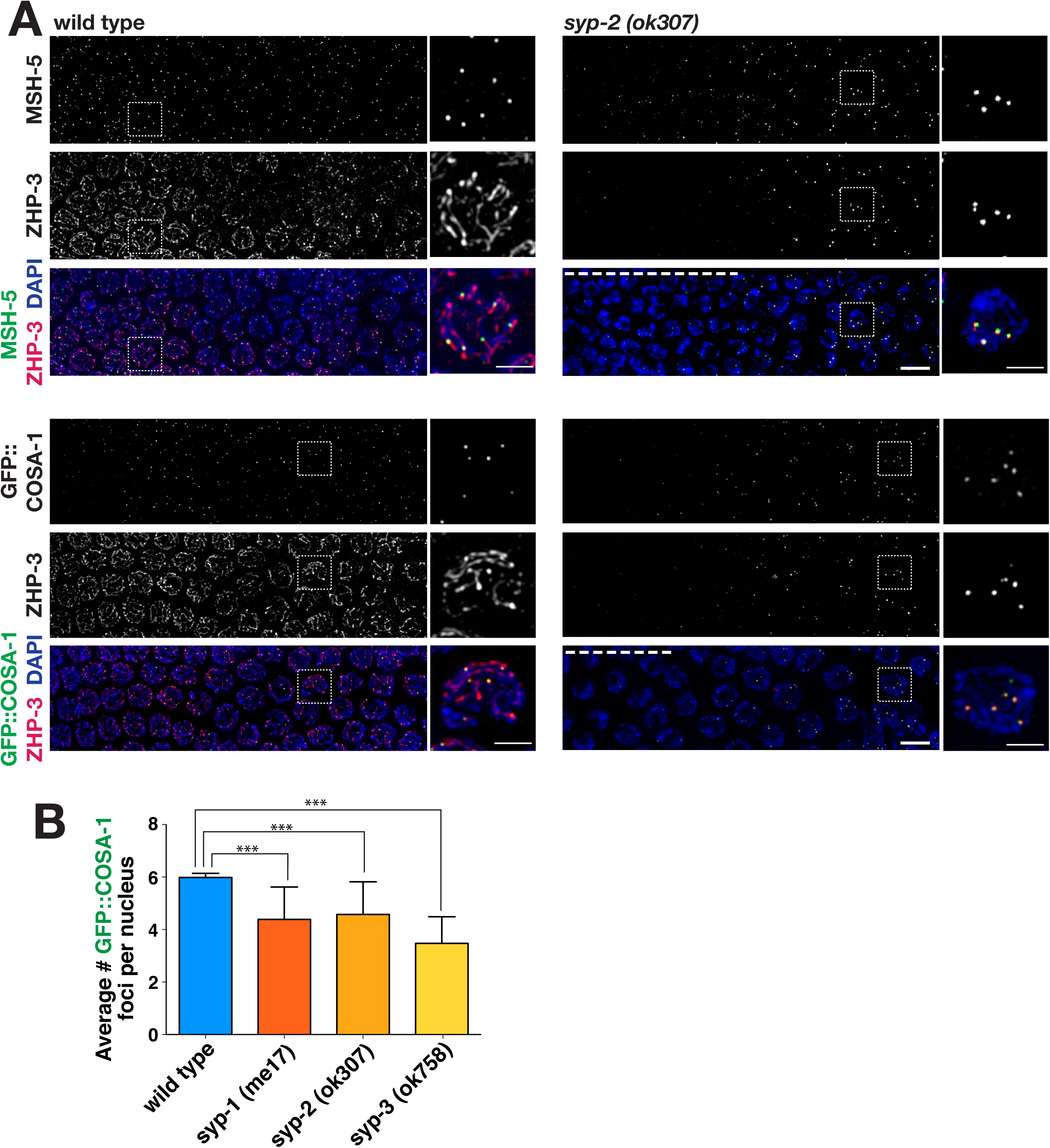
Co-localization of CO-promoting proteins to late pachytene foci in *syp* mutants. (A) Immunofluorescence images of mid-late pachytene region of germ lines from wild-type and *syp-2(ok307)* mutant worms, with meiotic prophase progressing from left to right. In wild-type nuclei at the mid-pachytene stage (left sides of wild-type panels), ZHP-3 is localized along the lengths of the chromosomes, MSH-5 is detected as foci in excess of the eventual number of COs, and COSA-1 foci are not detected. Upon transition to late pachytene, COSA-1 foci are detected at nascent CO sites, colocalized with MSH-5, and ZHP-3 tracks gradually reduce and retract toward the COSA-1 foci. In the *syp-2* mutant panels, nuclei at the left sides of the images exhibit characteristic DAPI signals reflecting prolonged persistence of chromosome clustering and chromosome movement (dashed white line). However, upon eventual release from chromosome clustering and transition to a late pachytene-like dispersed chromosome organization, ZHP-3 is detected as foci, most of which colocalize with MSH-5 (top) or GFP::COSA-1 (bottom). As the ZHP-3, MSH-5 and COSA-1 foci in the *syp* mutants are of weaker intensity than in wild-type, signal intensities in the *syp-2* images were boosted relative to controls to enable visualization of the foci. Dashed box indicates the nucleus that is enlarged in the adjacent image and scale bar on the enlarged images represents 2 µm. All other scale bar represents 5 μm. (B) Quantitation of GFP::COSA-1 foci in late pachytene nuclei for *syp* null mutants. Number of asterisks represent degree of statistical significance from a Mann Whitney test (***=*P*<0.0001). Error bars represent standard deviation. Number of nuclei scored for GFP::COSA-1: wild-type, n=505; *syp-1(me17)*, n=223; *syp-2(ok307)*, n=101; *syp-3(ok758)*, n=99.

The colocalization of pro-CO factors at multiple chromosome-associated sites in *syp* mutants is dependent on meiotic DSBs (Figure 2). Similar to the *spo-11* single mutant [which lacks endogenous DSBs; (Dernburg *et al.* 1998)], late pachytene nuclei in *spo-11; syp-1* and *spo-11; syp-2* double mutants typically have only occasional MSH-5 foci (0-1 foci per nucleus). This *spo-11*-dependent association of pro-CO factors to chromosome-associated foci in the absence of synapsis presumably reflects a proclivity of pro-CO factors to associate with each other and with the abnormal recombination intermediates present on the chromosomes in this context (Pattabiraman *et al.* 2017).

**Figure 2.**
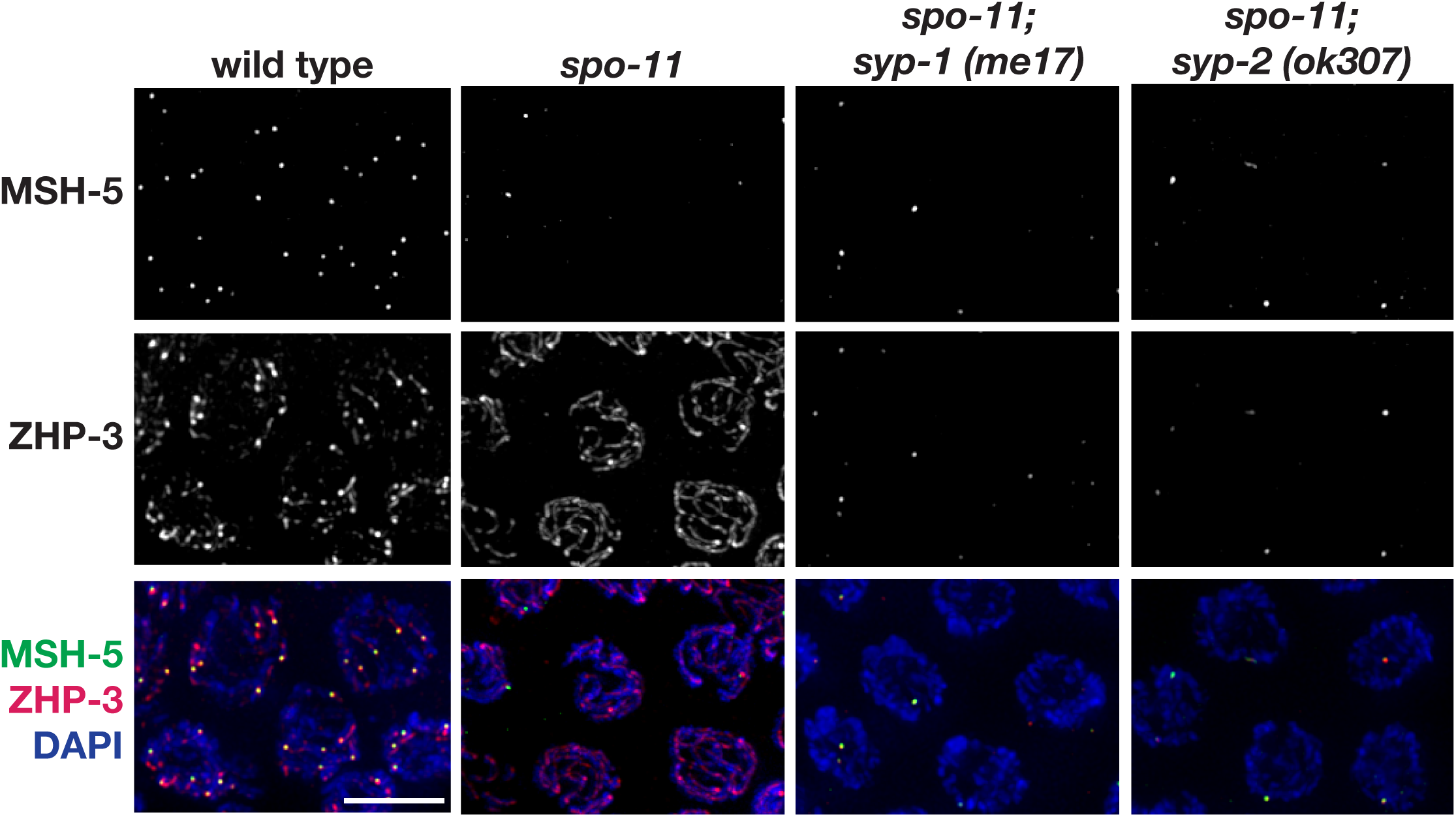
Co-localization of CO-promoting proteins in *syp* mutants is DSB dependent. Immunolocalization of MSH-5 and ZHP-3 in representative late pachytene nuclei from wild-type, *spo-11, spo-11; syp-1(me17)*, and *spo-11; syp-2(ok307)*. Similar to the *spo-*11 single mutant, late pachytene nuclei in *spo-11; syp-1* and *spo-11; syp-2* double mutants typically have only occasional MSH-5 focus (0-1), indicating that the presence of multiple foci in the *syp* single mutants is DSB dependent. In the *spo-11; syp* double mutants, most of the ZHP-3 detected is in foci colocalizing with MSH-5, in contrast to the *spo-11* single mutant in which ZHP-3 persists along the chromosome lengths. Scale bar represents 5 μm.

These observations in *syp* null mutants suggests that CO proteins have an inherent capacity to associate with DSB repair events, and that SYP proteins serve to constrain this activity both by limiting associations to appropriate interhomolog intermediates and by strengthening correct associations. Together, these findings indicate that meiotic chromosome structures collaborate to control the localization of CO-promoting proteins: 1) SYP proteins dictate and restrict where CO-promoting proteins can load; and, 2) correctly assembled chromosome axes restrict SYP proteins to load only between paired homologs. These features promote formation of interhomolog COs by ensuring that CO maturation occurs only in a productive manner, between properly aligned homologous chromosomes.

### CO-promoting proteins specifically associate with synapsed chromosome segments in mutants with limited synapsis

We examined localization of CO-promoting proteins and SYP-1 in late pachytene nuclei from worms homozygous for partial loss-of-function mutations affecting chromosome axis protein HIM-3 (Figure 3; Figure S1). In the *him-3(e1256)* mutant, in which partial impairment of chromosome axis function (Zetka *et al.* 1999) results in SYP-1 loading along only a subset of chromosome pairs (in most cases, four chromosomes), COSA-1 foci were associated only with the synapsed chromosomes, where extensive SYP-1 stretches were present (Figure 3A). Most SYP stretches in the *him-3(e1256)* mutant harbored only a single COSA-1 focus, and the average number of COSA-1 foci detected in late pachytene nuclei corresponded to the eventual number of chiasmata present in diakinesis-stage oocytes (Figure 3B; *P<*0.0001, Mann Whitney), indicating that the COSA-1 foci detected in this mutant represent *bona fide* CO events. COSA-1 foci were also invariably associated with the limited SYP-1 stretches present in the *him-3(me80)* mutant, which exhibits extensive synapsis only for the X chromosomes and only short stretches of synapsis on a subset of autosomes (Figure 3A). Similar to *him-3(e1256)*, we similarly detected only a single COSA-1 focus associated with any given SYP-1 stretch in *him-3(me80)* mutants. Thus, although our analysis of *syp* null mutants indicated that pro-CO factors do have a capacity to associate with DSB repair sites when SYP proteins are absent (Figures 1 and 2), our analysis of COSA-1 foci in the *him-3(e1256)* and *him-3(me80)* mutants suggests that when SYP proteins are present, they constrain where COSA-1 will localize (Figure 3). Additionally, in these mutants, SYP-1 loads in stretches prior to the COSA-1 localization, thereby supporting the conclusions with the *syp* mutants that the SYP proteins are directing where CO-promoting proteins can bind. Further, recent studies have shown that the SC-CR proteins are preferentially stabilized on chromosomes containing CO or CO-like events (Machovina *et al.* 2016; Nadarajan *et al.* 2016; Pattabiraman *et al.* 2017). Thus, the SC-CR proteins regulate the interaction of the CO-promoting proteins with the DSB site.

**Figure 3.**
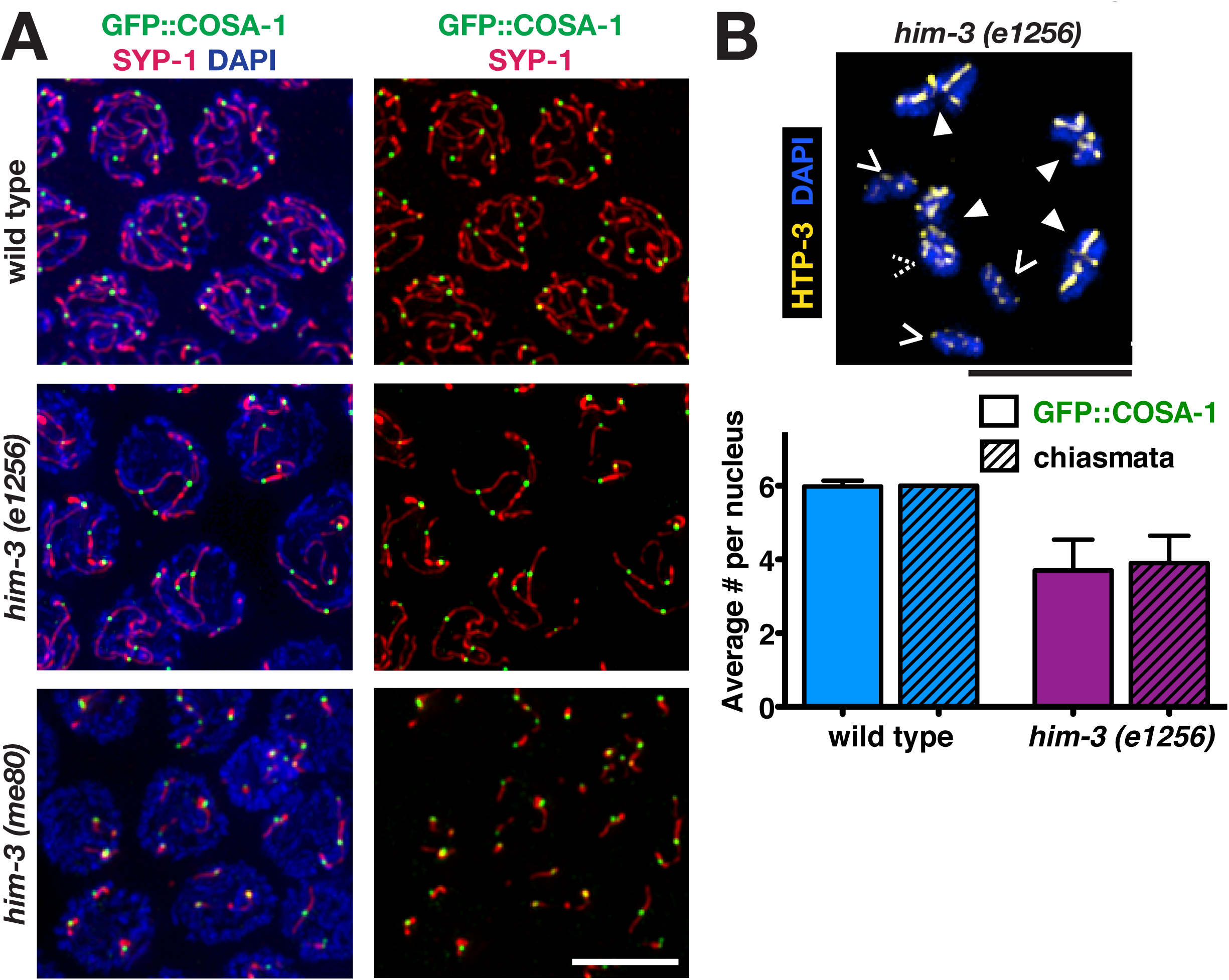
GFP:COSA-1 specifically associates with synapsed chromosome segments in mutants with limited synapsis. (A) Immunofluorescence images of representative nuclei in the late pachytene regions of germ lines from worms of the indicated genotypes, in which the SC central region protein SYP-1 localizes: along the lengths of paired homologs (wild-type); along the lengths of a subset of chromosomes (*him-3(e1256)* mutant); or in several short stretches (*him-3(me80)* mutant). Nearly all GFP::COSA-1 are associated with the chromosomes or chromosome segments where SYP-1 localized. (B) Top, immunofluorescence image of a representative diakinesis nucleus from the *him-3(e1256)* mutant, shown with DAPI (blue) and chromosome axis component HTP-3 (yellow) to visualize the chiasmata. Chiasmata were visualized and counted using 3D-rotations; solid arrowheads indicate bivalents connected by chiasmata, while carets indicate achiasmate chromosomes (univalent) (dashed caret indicates a univalent hidden in this projection). Bottom, bar graph depicting quantitation of GFP::COSA-1 foci in late pachytene nuclei and chiasmata in diakinesis nuclei for wild-type and the *him-3(e1256)* partial loss-of -function chromosome axis mutant; error bars indicate standard deviation (*P*<0.0001, Mann Whitney). Number of late pachytene nuclei scored for COSA-1 foci: wild-type, n=505; *him-3(e1256)*, n=161. Number of nuclei scored for chiasmata: wild-type, n=28; *him-3(e1256)*, n=40. All scale bars represent 5 µm.

Notably, previous studies of the *him-3(me80)* partial loss of function mutant have demonstrated that in addition to the formation of discontinuous SYP stretches in the *him-3(me80)* mutant, there are increases both in double COs by genetic assays and diakinesis bivalents with two chiasmata (Couteau *et al.* 2004; Nabeshima *et al.* 2004).

Based on our results demonstrating that each stretch of SYP in the *him-3(me80)* mutant usually receives a single COSA-1 focus along its length (Figure 3A) and that the number of COSA-1 foci correlates with the number of eventual chiasmata (Figure 3B), we provide further support to the original hypothesis posed that the occurrence of double COs and two-chiasmata diakinesis bivalents in *him3(me80)* mutant is due to discontinuous stretches SC assembly.

### Localization of CO-promoting proteins tracks with SYP-1 stretches when synapsis occurs incorrectly between sister chromatid pairs

We also examined localization of the CO-promoting proteins in a null mutant for *rec-8*, a component of one of the meiosis-specific cohesin complexes, which is required for pairing of the homologous chromosomes (Pasierbek *et al.* 2001; Severson *et al.* 2009). In contrast to wild-type where SYP-1 is loaded between paired homologous chromosomes, *rec-8(ok978)* null mutants inappropriately load SYP-1 between the 12 sister chromatid pairs (Figure 4A). When CO-promoting proteins were visualized in the *rec-8(ok978)* mutant, CO-promoting proteins were detected along the SYP-1 stretches in late pachytene nuclei (Figure 4B; Figure S1) and between condensed pairs of sister chromatids in diplotene and diakinesis phase nuclei (Figure 4D). Interestingly, 99% of *rec-8* mutants localized COSA-1 to no more than 12 sites in each late pachytene nucleus (Figure 4C). Further, ZHP-3 and MSH-5 similarly associate with COSA-1 foci strongly suggesting that recombination occurs between sister chromatids in this mutant (Figure S1).

**Figure 4.**
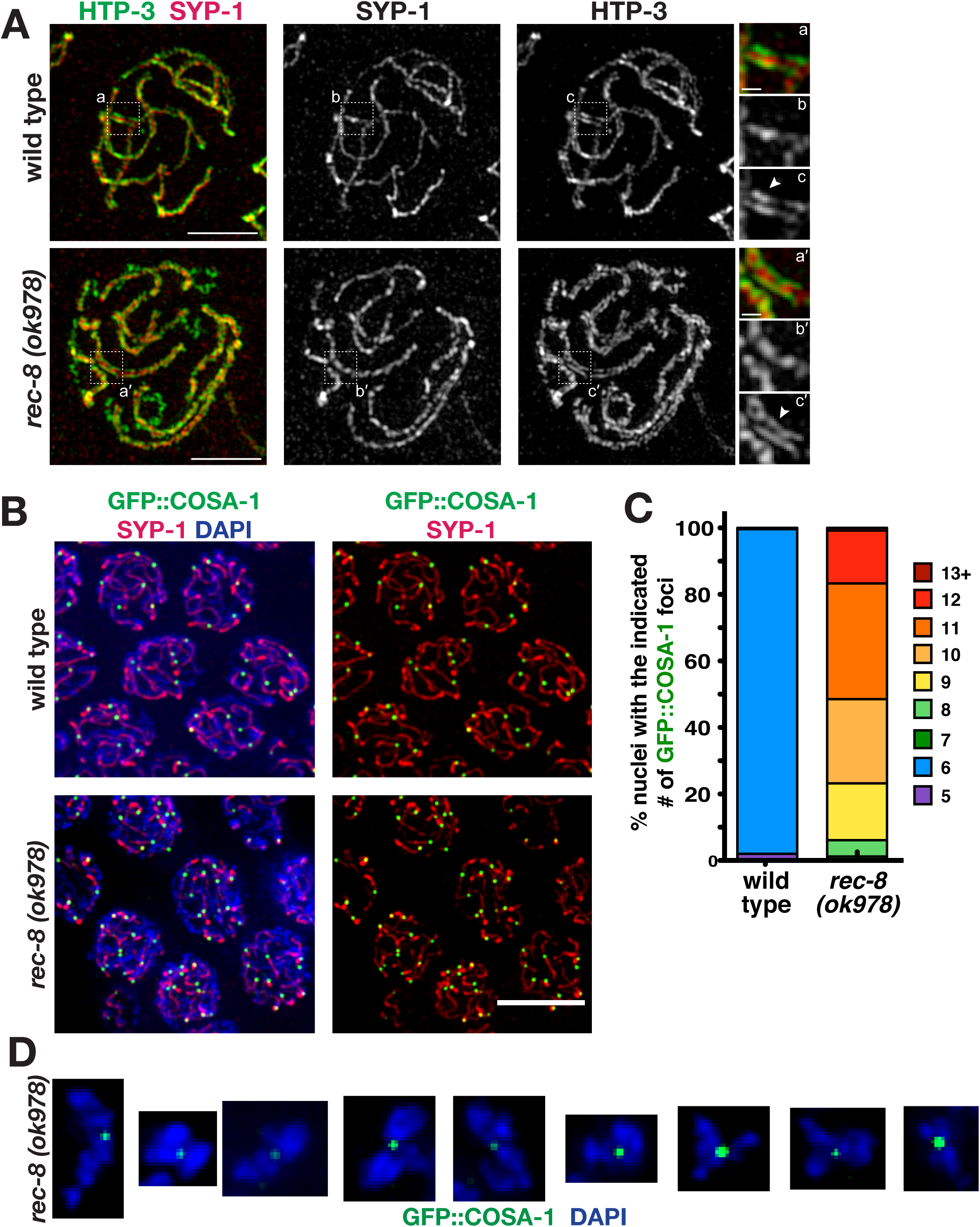
CO-promoting proteins associate with SYP-1 stretches in a *rec-8* mutant where synapsis occurs between sister chromatid pairs. (A) Structured illumination microscopy images of SYP-1 and chromosome axis component HTP-3 in representative pachytene nuclei, showing that SYP-1 localizes between pairs of HTP-3 tracks in both wild-type and *rec-8(ok978)* indicating that synapsis occurs between sister chromatid pairs in the *rec-8* mutant. White dash box indicates the enlarged region of SC depicted in the smaller images on the right. The arrowhead identifies a region where both lateral elements of the SC are visible indicated by the two tracks of HTP-3. Scale bars for whole nucleus images represent 2 µm and scale bars for smaller enlarged SC segments represents 250 nm. (B) Immunolocalization of SYP-1 and GFP::COSA-1 in fields of nuclei from the late pachytene regions of wild-type and *rec-8(ok978)* germ lines. In the *rec-8* mutant, SYP-1 localizes along the lengths of unpaired chromosomes, and COSA-1 foci are much more abundant than in wild-type. Scale bar represent 5 µm. (C) Stacked bar graph showing percentages of nuclei with indicated numbers of GFP::COSA-1 foci in wild-type and *rec-8(ok978).* Number of late pachytene nuclei scored for COSA-1 foci: wild-type, n=505; *rec-8(ok978)*, n=245. (C) Immunolocalization of GFP::COSA-1 in DAPI stained diakinesis bivalents from *rec-8(ok978)* germ lines. In the *rec-8* mutant, GFP::COSA-1 localizes between the two sister chromatids.

The numbers we observed for COSA-1 foci formation in *rec-8(ok978)* null mutants is consistent with interference occurring along synapsed sister chromatid pairs. As there are 12 pairs of sister chromatids in the *rec-8(ok978)* null mutants and we only very rarely observe greater than 12 COSA-1 foci (0.8% of all nuclei), the average number of 10.4 ± 1.2 COSA-1 foci per nucleus observed in *rec-8(ok978)* null mutants reflects an imposed limitation of COSA-1 foci by the number of pairs of sister chromatids. As there are regions of occasional non-homologous synapsis, those nuclei exhibiting less than 12 COSA-1 foci most likely have a sister chromatid pairs with regions of non-homologous synapsis. Given our previous study demonstrating a role for the SC in promoting CO interference (Libuda *et al.* 2013), in these cases of non-homologous synapsis within the *rec-8(ok978)* null mutant, it is possible that these partially synapsed sister chromatid pairs are being recognized as a signal “module” or chromosome. Hence, interference, which is occurring along one set of sisters, may be transmitted along the other set of sister chromatids to which the pair is partially synapsed. Overall, these results further support the hypothesis that the SYPs may serve as the scaffolding along which a signal may be propagated in *C. elegans*.

### Connection of the sister chromatids at diakinesis in a rec-8 null mutant is dependent on CO-promoting proteins

The localization of COSA-1 foci between condensed sister chromatid pairs during diakinesis in *rec-8* null mutants (Figure 4D) suggests a potential role for COSA-1 in holding together sister chromatids. Moreover, the presence of COSA-1 between the sisters at diakinesis in *rec-8* mutants suggests that COSA-1 is marking a DSB-dependent event. Previous studies have shown that *rec-8* null mutants will frequently equationally separate the sister chromatids at the first meiotic division and that the sister chromatids are held together in a DSB-dependent manner prior to the first meiotic division (Severson *et al.* 2009; Severson and meyer 2014). Thus, it is possible that the COSA-1 foci are marking intersister COs, which are being used to equationally separate the sister chromatids at meiosis I.

To determine whether the localization of COSA-1 between condensed sister chromatids in diplotene and diakinesis nuclei of the *rec-8* null mutant was required to hold sister chromatid together, we assessed the number of diakinesis bivalents in *rec-8, cosa-1*, and *cosa-1; rec-8* double mutant (Figures 5A and 5B). Similar to previous studies, we found that both *rec-8* and *cosa-1* null mutants displayed on average twice as many DAPI-staining bodies at diakinesis as wild-type indicating that the CO connections between the homologs are absent resulting in the formation of univalents (Severson *et al.* 2009; Yokoo *et al.* 2012). Similar to Crawley *et al.*, we found that the *cosa-1; rec-8* double mutant displayed on average 18 DAPI-staining bodies at diakinesis (and occasionally 24), suggesting that in *rec-8* mutants the sister chromatids are being held together in a COSA-1 dependent manner (Crawley *et al.* 2016). To determine if the SC-CR is responsible for the COSA-1 localization and maintaining the intersister connections in diakinesis, we assessed the number of DAPI-staining bodies at diakinesis in a *rec-8; syp-2* double mutant. Surprisingly, rather than phenocopying *cosa-1; rec-8*, the *rec-8; syp-2* double mutant occasionally displayed severe chromosome fragmentation defects indicative of DSB repair defective mutants (Figure 5A) (Colaiacovo *et al.* 2003; Crawley *et al.* 2016). Thus, the SC-CR is critical in *rec-8* mutants to promote the formation of COSA-1 foci, which function in maintaining intersister connections possibly through the formation of intersister crossovers.

**Figure 5.**
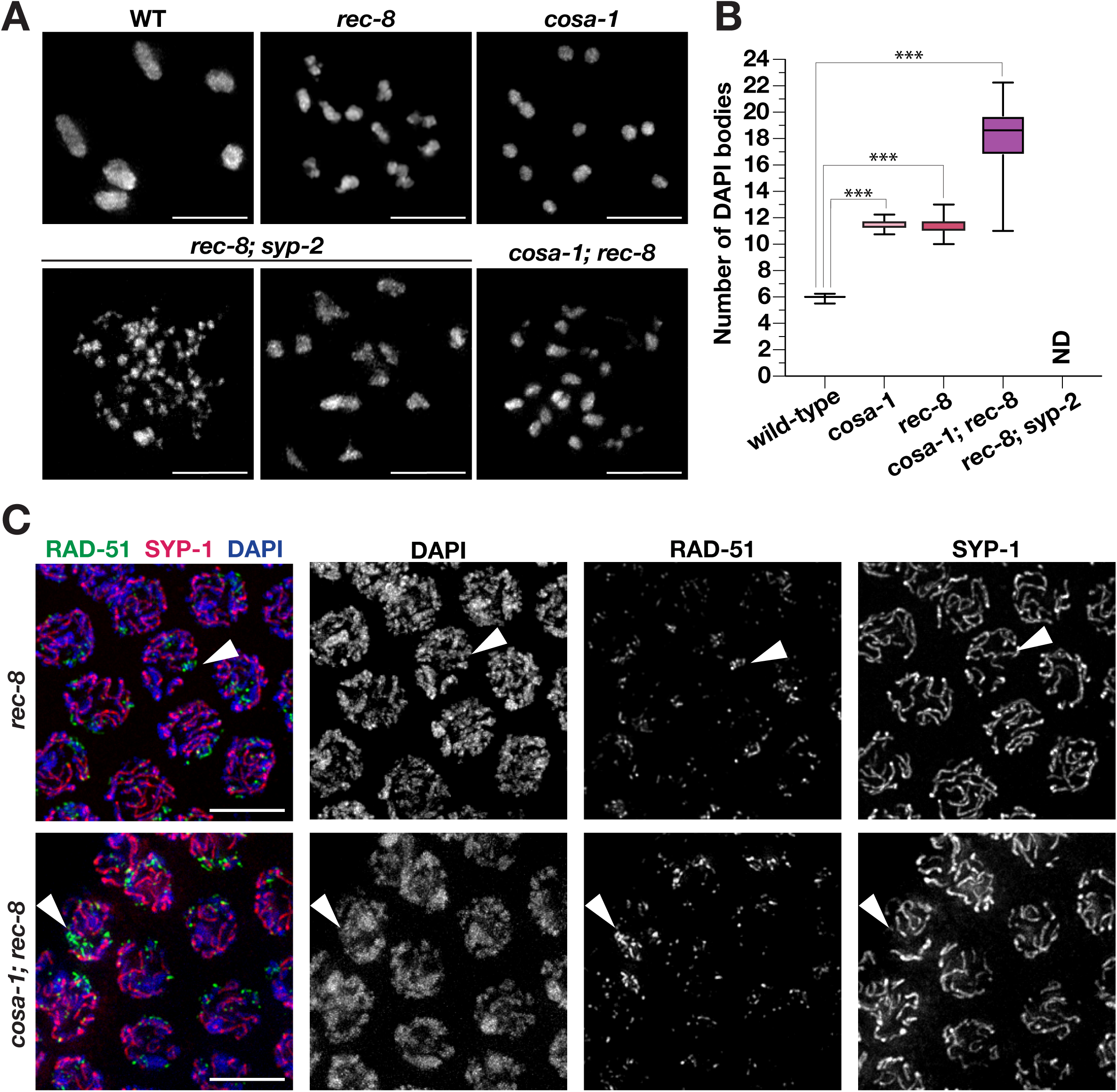
Connection of sister chromatids during diakinesis is dependent on CO-promoting proteins. (A) Representative image of DAPI stained nuclei from the diakinesis region of wild-type, *rec-8*, *cosa-1*, *cosa-1; rec-8*, and *rec-8; syp-2* germ lines. Both *rec-8* and *cosa-1* mutant are defective in homolog pairing, while the *cosa-1; rec-8* double mutant is also unable to maintain sister chromatid pairing. The *rec-8; syp-2* double mutant has severe chromosome fragmentation likely caused by an inability to associate with a partner to repair DSBs. Scale bars represent 5 µm. (B) Box plot depicting the quantification of the DAPI bodies in panel A. Number of nuclei scored for DAPI bodies: wild-type, n=213; *rec-8*, n=193; *cosa-1*, n=226; *cosa-1;rec-8*, n=162. Due to the degree of chromosome fragmentation in *rec-8; syp-2* these nuclei could not be accurately scored for DAPI bodies (ND = not determined). Number of asterisks represent degree of statistical significance from a Mann Whitney test (***=*P*<0.0001). (C) Immunolocalization of RAD-51 and SYP-1 in the late pachytene regions of *rec-8(ok978)* and *cosa-1; rec-8(ok978)* germ lines. RAD-51 forms concentrated numbers of foci in regions where SYP-1 is absent. Scale bars represent 5 µm.

The striking chromosome fragmentation defects in the *rec-8; syp-2* double mutant suggest that the chromosomes may need to be associated with a partner to initiate DSB repair. Since we were able to observe regions of desynapsis (Figure 4A) in *rec-8* mutants, we examined whether DSBs accumulate in these regions. While the initial repair of DSBs by the loading and off-loading of RAD-51 looks similar between *rec-8* and *cosa-1; rec-8* mutants (Figure S2), RAD-51 accumulated in both mutants on chromosome stretches where SYP-1 had not loaded (Figure 5C). While the percentage of nuclei without SYP-1 do not correspond to the percentage of nuclei with fragments, it is possible that we are not able to detect all the chromosome fragments with DAPI. Thus, these data suggest that DSB repair requires the SC-CR proteins to promote partner association. In the absence of the SC-CR, cohesion remains intact and likely enables enough partner association to allow for DSB repair with the sister chromatid.

However, in the absence of both the SC-CR and cohesion partner associations, DSBs are unable to repair resulting in chromosome fragmentation.

### CO-promoting proteins associate with SYPs even in the context of aggregates that form in null mutants lacking meiotic chromosome axis components

In addition to CO-promoting proteins preferentially associating with SYP stretches in the context of limited or inappropriate synapsis, we also found that CO-promoting proteins associate with SYP-1 even when SYPs are unable to load onto chromosomes, instead forming aggregates (Figure 6; Figure S1). In null mutants for the lateral elements HTP-3 and HIM-3, SYP proteins are unable to load onto chromosomes and instead form an aggregate within the nucleus (Couteau *et al.* 2004; Goodyer *et al.* 2008). Specifically, in the *him-3* null mutant, late pachytene nuclei typically contain a single elongated SYP-1 aggregate (Figure 6A). In the *htp-3* null mutant, late pachytene nuclei contain one or sometimes two aggregates per nucleus (Figure 6A). In both of these null mutants for *him-3* and *htp-3*, we observe CO-promoting proteins associated with the SYP aggregate (Figure 6A; Figure S1). Further, in most cases, only a single COSA-1 focus is associated within a given SYP-1 aggregate (Figure 6A). Additionally, recent data show that the pro-CO factors ZHP-3 and COSA-1 also localize to these SC aggregates as a single focus (Rog *et al.* 2017), suggesting that COSA-1 has a strong tendency to associate with SYP-1 and that the ability to limit COSA-1 foci to a single site on a given SYP-1 structure is retained even when the SYP proteins are concentrated in a non-chromosomal location.

**Figure 6.**
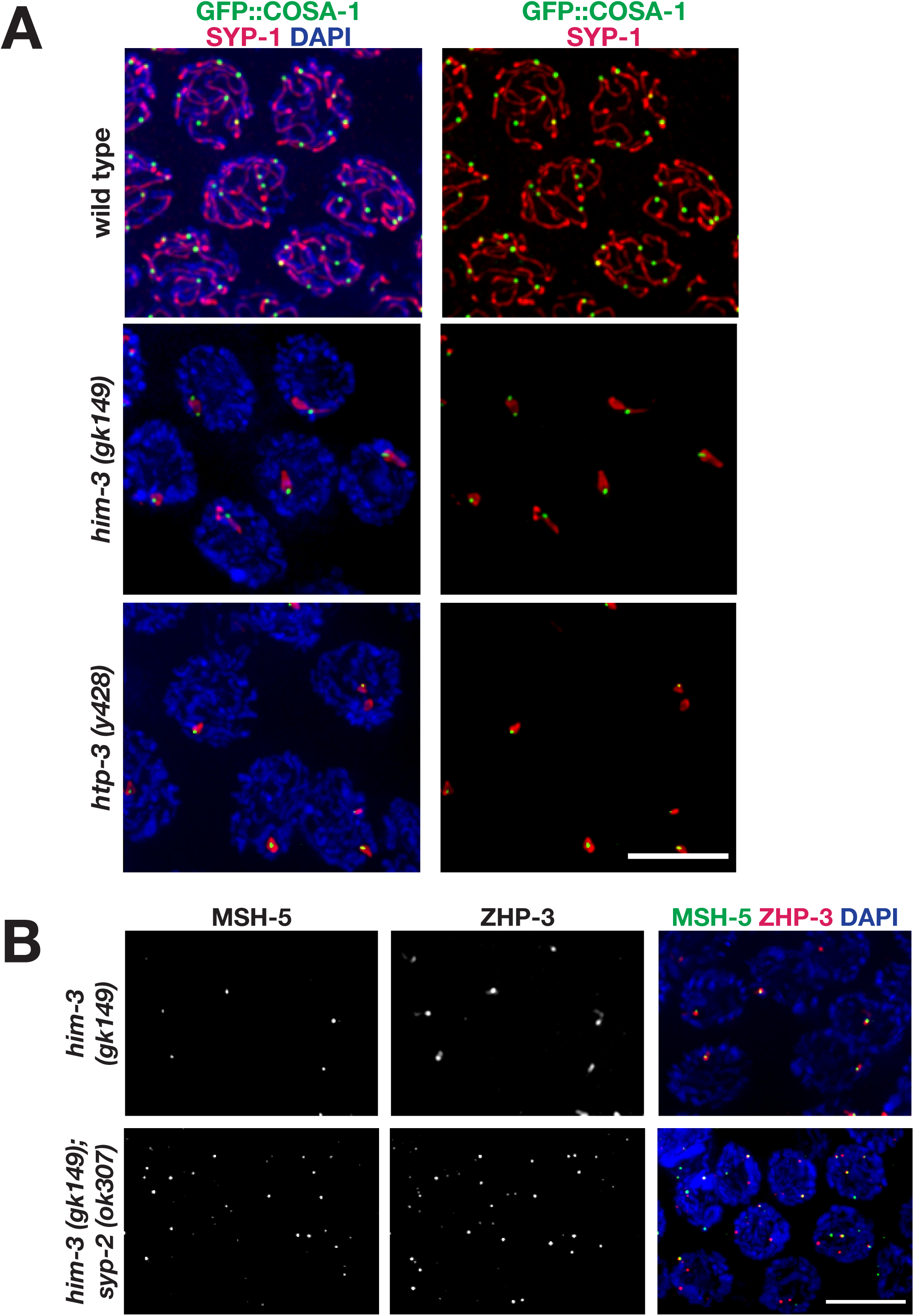
CO-promoting proteins associate with abnormal SYP-1 structures in mutants lacking meiotic chromosome axis components. Immunolocalization of SYP-1 and GFP::COSA-1 in nuclei from the late pachytene regions of null mutants lacking chromosome axis components HIM-3 or HTP-3. SC assembly is severely impaired in both the *him-3(gk149)* and *htp-3(y428)* null mutants, and SYP proteins instead assemble into abnormal aggregates known as polycomplexes. GFP:COSA-1 localization is consistently associated with these abnormal SYP-1 structures in both mutants. (B) Whereas MSH-5 and ZHP-3 are detected together at a single site per nucleus in the *him-3(gk149)* mutant, multiple foci are detected in nuclei in the *him-3; syp-2* double mutant (as in the *syp-2* single mutant). All scale bars represent 5 µm.

In the context of these meiotic chromosome axis mutants, the restriction of CO-promoting proteins to a single site per nucleus is dependent on the SYP proteins. In contrast to the *him-3(gk149)* null mutant, we found that in the *him-3; syp-2* double mutant, CO-promoting proteins localize to multiple chromosome-associated sites in late pachytene nuclei (Figure 6B), similar to the *syp-2* single mutant (Figure 1). This finding suggests that aggregation of SYP proteins to a single site preferentially stabilizes the association of CO-promoting factors and constrains them to colocalize together with SYP-1 in a single compartment. When this constraint is released, the CO-promoting factors are free to localize together at DSB-dependent chromosomal sites.

### Partial depletion of SYP proteins results in an increased number of COSA-1 foci along a SYP-1 stretch

In previous work, we showed that partial depletion of the SYP proteins results in an increase number of COSA-1 foci and attenuates CO interference, implicating the SC central region in limiting and constraining CO numbers (Libuda *et al.* 2013). In our current work, we found that partial depletion of SYP-1 similarly resulted in increased occurrence of SYP-1 stretches with greater than 2 COSA-1 foci in the *him-3(e1256)* partial loss of function mutants (Figure 7). In 14% of control RNAi treated *him-3(e1256)* nuclei, we observe two COSA-1 foci along a SYP-1 stretch. Upon *syp-1* partial RNAi treatment, the number of SYP-1 stretches with greater than two COSA-1 foci significantly increases to 41%. Notably, only continuous stretches of SYP-1 were counted, therefore if a single chromosome had two discontinuous stretches of SYP-1, then it was counted as two separated stretches of SYP-1. Quantitation of COSA-1 foci along entire individual chromosomes was not possible as the asynapsed chromosomes in *him-3(e1256)* made resolving individual chromosomes difficult. As several chromosomes appeared to have discontinuous stretches of SYP-1 upon *syp-1* partial RNAi, 41% is likely an under-estimate of the actual number of COSA-1 foci along a single chromosome. Further, partial depletion of SYP-1 in the *rec-8(ok978)* null mutant background also resulted in the frequent occurrence of SYP-1 stretches harboring two COSA-1 foci, even while reducing the fraction of chromosomes associated with SYP-1 stretches (Figure 7). Together, our data reinforce the suggestion that the SC central region functions in regulating CO numbers and distribution.

**Figure 7.**
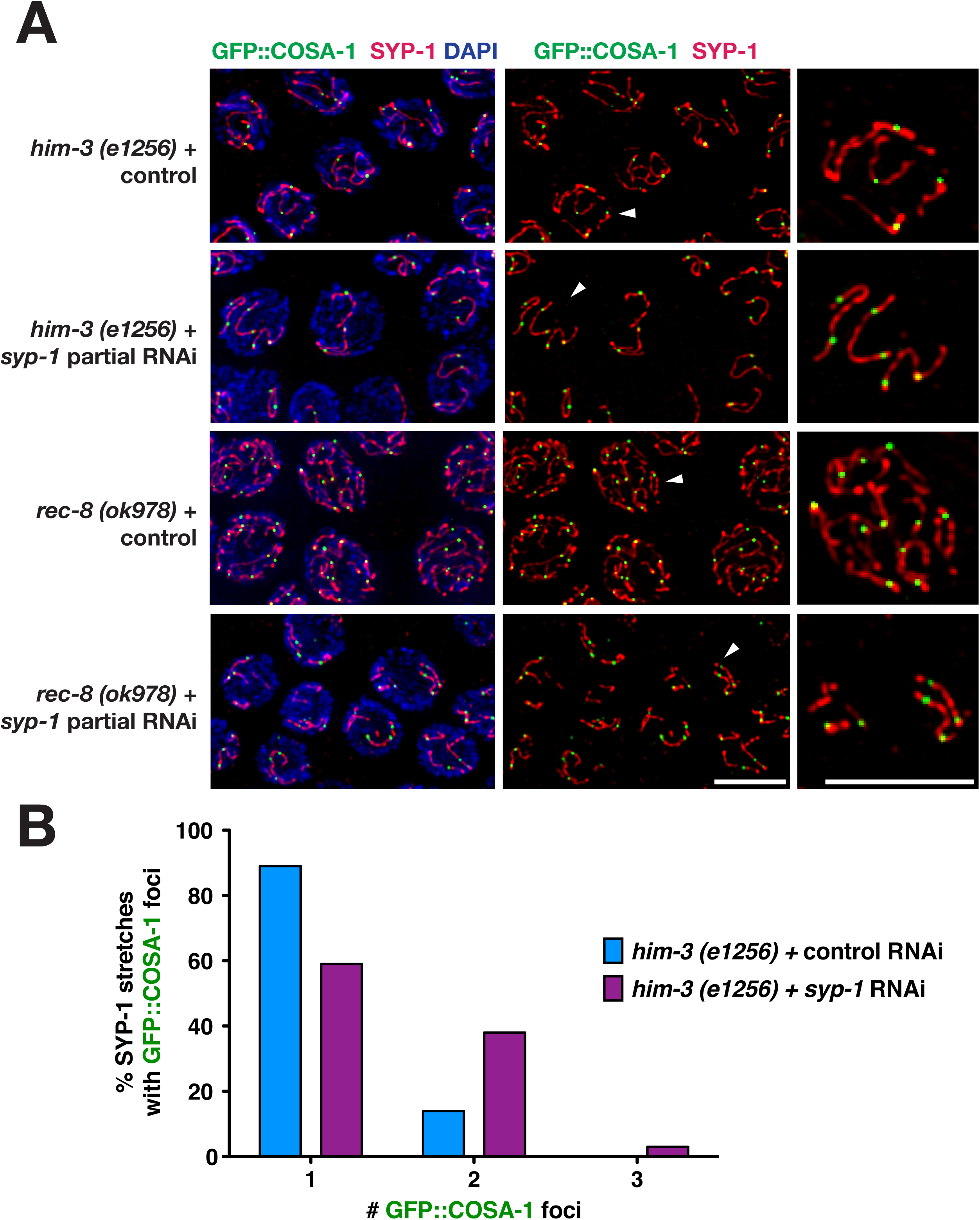
Increased numbers of COSA-1 foci along SYP-1 stretches following partial depletion of SYP-1. (A) Immunolocalization of GFP::COSA-1 and SYP-1 in late pachytene nuclei from control *him-3(e1256)* or *rec-8(ok978)* worms and from *him-3(e1256)* or *rec-8(ok978)* worm treated with *syp-1* partial RNAi. Arrowheads point to nuclei that are enlarged in panels on the right. As SYP-1 immunofluorescence intensities following *syp-1* partial RNAi often appeared weaker relative to controls, SYP-1 signal intensities were boosted for the *syp-1* partial RNAi panels to facilitate visualization of SYP-1 tracks (Libuda *et al.* 2013). Whereas SYP-1 is localized along most sister chromatid pairs in *rec-8* control nuclei, SYP-1 is detected along only a subset of sister chromatid pairs in the *rec-8; syp-1* partial RNAi nuclei. Further, in contrast to controls, nuclei from either *him-3(e1256)* or *rec-8(ok978)* mutants treated with *syp-1* partial RNAi often contained SYP-1 stretches with SYP-1 stretches harboring more than one GFP::COSA-1. Scale bars represent 5 µm. (B) Graph showing quantitation of the percentage of contiguous SYP-1 stretches in control *him-3(e1256)* nuclei (n=103) or *him-3(e1256)* nuclei treated with *syp-1* partial RNAi (n=100) that had the indicated numbers of GFP::COSA-1 foci.

## Concluding remarks

The use of immunofluorescence of the CO-promoting proteins in various meiotic chromosome structure mutants allowed us to visualize how various components of the SC influence CO-promoting protein localization and function. Our data indicate that that CO-promoting proteins have a capacity to localize to DSB-dependent events outside of the context of the SC, but that this inherent affinity for DSB events is constrained by the location of the subunits of the SC central region (SYP proteins). How the SC central region proteins physically interact at the molecular level with the CO-promoting proteins to direct their localization may be determined by futures studies investigating the structure of these proteins.

## Acknowledgements

We thank A. Dernburg, A. Villeneuve, and M. Zetka for antibodies and the CGC (funded by National Institutes of Health (NIH) P40 OD010440), B. Meyer, and A. Villeneuve for strains. We thank A. Villeneuve and members of the Libuda Lab, especially N. Kurhanewicz and E. Toraason, for comments on the manuscript. This work was supported by the National Institutes of Health R00HD076165 and R35GM128890 to DEL. DEL is also a Searle Scholar and recipient of a March of Dimes Basil O’Connor Starter Scholar award.

---

**Figure S1.**
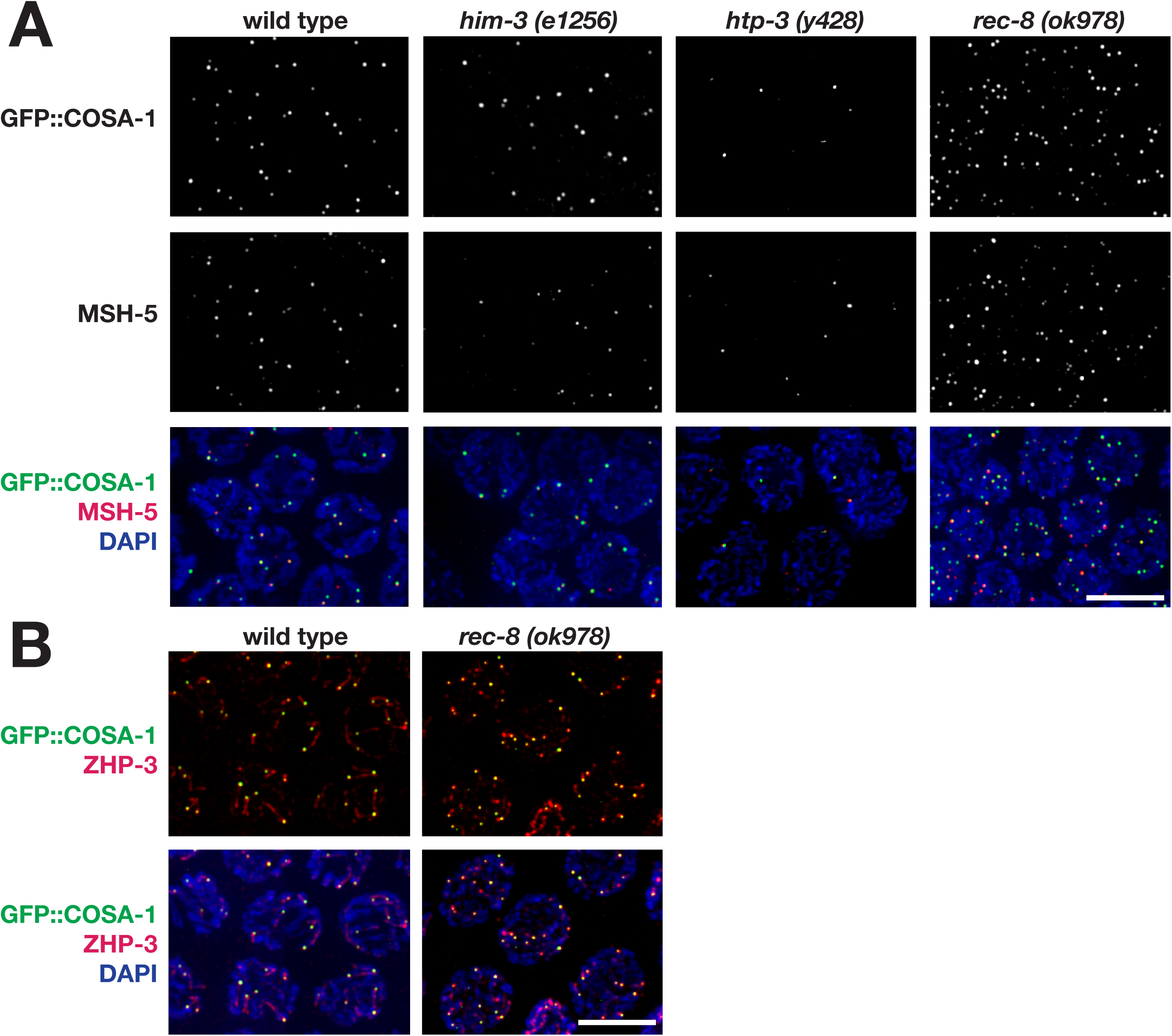
Co-localization of CO-promoting proteins in *him-3, htp-3*, and *rec-8* mutants. (A) Immunolocalization of GFP::COSA-1 and MSH-5 from the late pachytene regions of wild-type, *him-3(e1256), htp-3(y428)* and *rec-8(ok978)* germ lines. (B) Immunolocalization of GFP::COSA-1 and ZHP-3 from the late pachytene regions of wild-type and *rec-8(ok978)* germ lines. Scale bar represent 5 µm.

**Figure S2.**
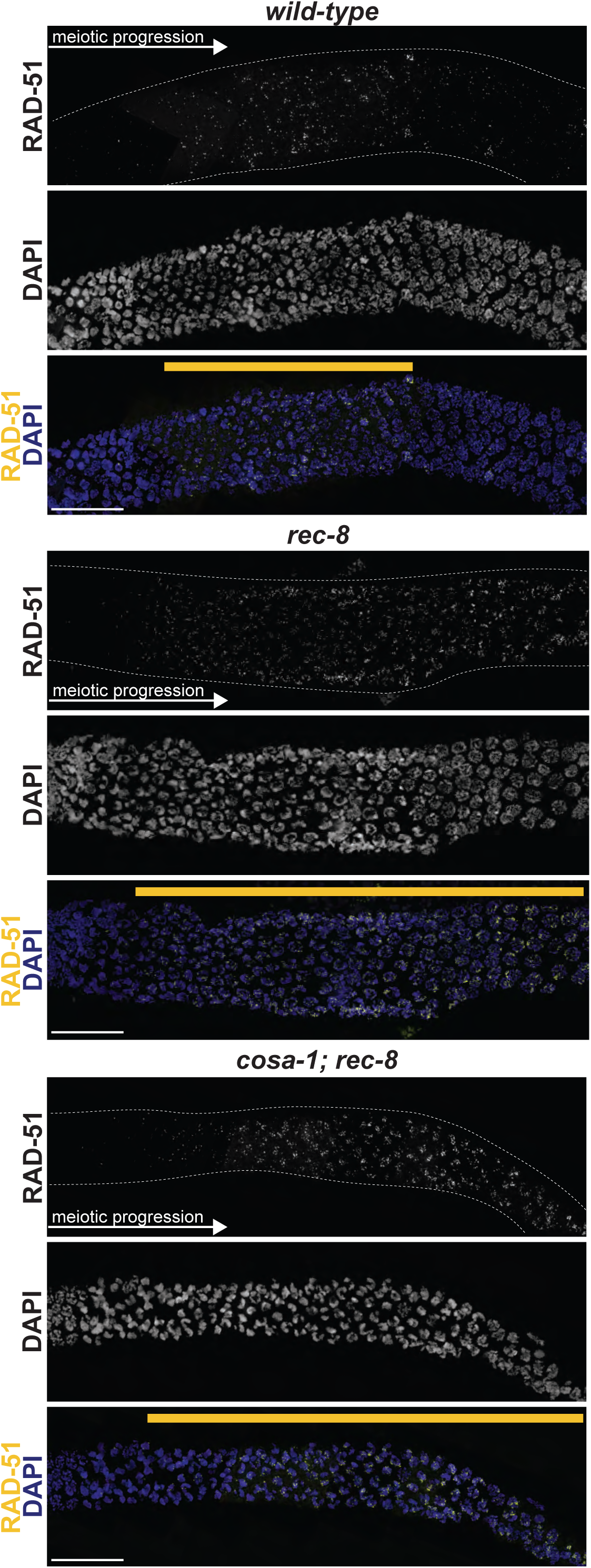
RAD-51 localization in *rec-8* and *cosa-1; rec-8*. Immunolocalization of RAD-51 in whole gonads from wild-type, *rec-8*, and *cosa-1; rec-8*. Both *rec-8* and *cosa-1; rec-8* have a persistence of RAD-51 foci. Yellow line indicates the length of the RAD-51 zone in each genotype. White-dashed line shows the outline of the gonad. Scale bar represents 20 µm.

